# Defence systems drive accessory genome interactions in *Pseudomonas aeruginosa*

**DOI:** 10.1101/2025.05.06.652404

**Authors:** Charlotte E. Chong, Aaron Weimann, Aleksei Agapov, Joanne L. Fothergill, Michael A. Brockhurst, Julian Parkhill, R. Andres Floto, Mark D. Szczelkun, Edze R. Westra, Multi-Defence Consortium, Kate S. Baker

## Abstract

Bacterial genomes represent dynamic ecological systems in which mutational selection and dynamic accessory genome element complements drive evolution. Emerging evidence suggests that bacterial defence systems, which protect against phages and other mobile genetic elements, interact through cooperative, competitive, and antagonistic relationships. These interactions can influence horizontal gene transfer, shape phage susceptibility, and the diversification of genome composition across environments. Recent ecological studies describe non-random co-occurrence and avoidance of defence systems, suggesting that they may emerge from ecological and evolutionary interactions, rather than by chance. This necessitates exploring whether these patterns reflect functional compatibility, shared selection pressures imposed by ecological niche, or if they merely arise from co-localisations of convenience. Understanding these patterns is key to elucidating how the accessory genome evolves and how defence systems constrain or facilitate genome plasticity. To investigate these patterns, in their genomic context, and provide a resource for future investigations, we analysed the distributions of defence systems and other accessory genome elements in a recently curated global dataset of 2,940 *Pseudomonas aeruginosa* genomes. Genomic defence system content varied by ecological niche, with higher numbers per genome in non-cystic fibrosis derived isolates (average n=7.9) compared to cystic fibrosis-derived isolates (average n=6.5). We identified multiple associations (n=426) and dissociations (n=50) among defence systems, and among other accessory genome elements, many of which had a plausible biological explanation. We also quantitated the relative interactions among accessory genome elements which revealed that defence systems and anti-defence systems engage in the most accessory genome interactions, compared with e.g. antimicrobial resistance genes, plasmids, phages, suggesting that systems are a major driving force in the ecological dynamics of the bacterial accessory genome. Together, these patterns provide new insights into the evolutionary forces shaping bacterial genomes, provide a valuable resource of quantitated interactions in a highly curated *Pseudomonas aeruginosa* dataset, and establish a framework for future mechanistic and ecological investigations of defence system interactions.

## Introduction

Bacterial evolution is partially driven by the acquisition of beneficial traits like antimicrobial resistance (AMR), often facilitated by horizontal gene transfer (HGT) of mobile genetic elements (MGEs) such as plasmids, integrative conjugative elements (ICEs) and phages (1, 2). In this study, we define these collectively as ‘Accessory Genome Elements (AGEs)’, encompassing both MGEs (including phage, plasmids and ICEs) and functionally defined elements (such as defence systems (DSs), antimicrobial resistance genes (ARGs) and anti-defence systems (Table 1). These AGEs collectively contribute to bacterial adaptability, genome plasticity, and evolution but their can also affect fitness and, in some cases, cause cell death. In response to this evolutionary pressure, bacteria have developed immune responses against extraneous AGEs. Bacteria can contain many complex defence systems (DSs) to defend against harmful invasion ensuring a balance between adaptability and genome integrity (3). DSs such as restriction-modification (RM) and CRISPR-Cas are powerful contributors to the control of HGT but in some case can stabilise AGEs like phage and plasmids (4, 5).

**Table 1.** Overview of accessory genome elements (AGEs) considered in this study.

| Category | Type | Examples | Functional role |
| --- | --- | --- | --- |
| **Mobile element** | Phage | Temperate and lytic phages- Dobby, fMGyn-Pae01 | Mediate HGT, impose selection for DSs |
|  | Plasmid | Conjugative and non-conjugative plasmids- IncQ2 type plasmids | Transfer genes, including ARGs and DSs |
|  | Integrative conjugative elements (ICEs) | Tn4371-like elements, bph-sal | Mobilise and integrate genetic material between bacterial genomes |
| **Functional element** | Defence systems (DSs) | Restriction modification (RM), CBASS, AbiE, gabija | Protect against phages and MGEs |
|  | Antimicrobial resistance genes (ARGs) | blaOXA, sul1, ermB | Confer resistance to antimicrobials |
|  | Anti-defence systems | Anti-CRISPRs (Arcs) and Anti-CBASS | Inhibit defence systems |

Multiple and diverse DSs are often found within bacterial genomes, with an average of five DSs in over 20,000 sequenced prokaryotic genomes (6, 7). Furthermore, recent studies demonstrate that DSs may be found clustered in defence islands (3, 7–11) which are commonly associated with mobilisable AGEs, and may facilitate their transfer between genomes (12, 13). The proximity to mobile AGEs and resultant genome plasticity offers a fitness benefit to bacteria living and responding to phage-rich environments (3). It has been hypothesised that defence islands form within bacterial genomes due to mechanistic synergism between clustered DSs promoting co-localisation and mobilisation between genomes (11, 14). Such aggregation is consistent with that seen for AMR and pathogenicity genes on mobile AGEs, such as plasmids, ICEs and phage (11, 15, 16), suggesting that defence islands may be similarly promoted by evolutionary forces including HGT, shared selective advantage, and genetic linkage. Owing to these multiple influences, it remains challenging to identify those DS interactions which represent putative mechanistic interactions, and those which result from chance co-localisation.

To address these deficiencies in understanding, here we comprehensively analyse interactions among DSs, and other AGEs, including plasmids, ICEs, ARGs, phages, and the newly described anti-defence systems in a recently curated global dataset of 2,940 *P. aeruginosa* genomes (17). *P. aeruginosa* is an ideal organism to study DS interaction as previous studies have found it contains varied DSs (6, 9, 18). It is a widely distributed, Gram-negative bacterium and is found in a diverse range of ecological niches. As an opportunistic pathogen, it is a major concern in hospital settings, causing a broad range of infections, particularly in individuals with cystic fibrosis (CF). Its adaptability has contributed to its status as a World Health Organisation high-priority AMR pathogen (19). Furthermore, DSs were recently revealed to play a key role in its adaptation to different transmission models, making it a valuable model for studying bacterial immunity (9, 17, 20). We identify a weak but significant effect of niche (CF vs. non-CF) in driving accessory genome complements and then go on to characterise ecological interactions among DSs and a wide range of AGEs, recovering multiple putative phylogenetically independent interactions for downstream mechanistic investigation. We also demonstrate that DSs and anti-defence systems are the principal drivers of AGEs in *Pseudomonas* genomes providing valuable insights into microbial ecology, evolution, and potential mechanistic interactions.

## Methods

### Genomic dataset

*P*. *aeruginosa* genomes used in this study were previously described (17). We utilised 2,940 (from a total of 9,829 genomes) genomes from studies of AMR in healthcare settings (21–24), individuals with cystic fibrosis (17, 25, 26), bacteraemia (27) and non-CF bronchiectasis (28), from studies into high-risk *P. aeruginosa* (29–31) and from the International *Pseudomonas* Consortium (32) (Supplementary Table 1). Isolate genomes from 9 deeply sampled CF patients from the TeleCF study were (17) down sampled, and only genomes representative of diversity were included in the analysis. Specifically, only patient representative strains were retained (i.e. the oldest sample for each sequence within a patient) to avoid overrepresentation of clones from deeply sampled patients. Genomes were assembled from short-read sequence data, using the Velvet (33) or SPAdes (34) assembler using the Assembly Improvement pipeline (35), as described by Weimann, et al. (2024). Bacterial genomes were annotated with PROKKA v1.14.6 (36).

### Detection of accessory genome elements in bacterial genomes

DSs and anti-defence systems were detected in *P. aeruginosa* draft assembly FASTA files using DefenseFinder-DB v1.2.2 (accessed April 2023) and Anti-DefenseFinder-DB v1.3.0 (accessed December 2024) respectively (6). These databases contained 152 DSs and 41 anti-defence systems. Integrative conjugative elements (ICEs) and plasmids were detected in bacterial genomes using ICEberg v.2.0 (1032 database entries) (37) and PlasmidFinder v.2.1.6 (116 database entries) (38) databases via ABRicate v.1.0.1(39), using default parameters. Phage genomes present within the *P. aeruginosa* isolates were detected using geNomad v. 1.7.0 (40) and further characterised with PHASTEST webserver (∼410,000 viral sequences) (41). Additionally, ARGs were identified using NCBI AMRFinderPlus v. v3.11.11, which contains 6,428 ARG entries (42).

### Identifying co-occurrence

Coinfinder v.1.10 (43) was used for detecting significant interactions (i.e. associations and dissociations) among AGEs in the *P. aeruginosa* dataset, using the presence and absence data of defence systems, phage, plasmids, ICEs and ARGs, along with a maximum likelihood tree phylogenetic tree generated by Weimann *et al*, as input (Supplementary Table 1, see Figure 1C in Weimann, et al 2024) (17). This tree was constructed using FastTree v2.1.10 (44) based on a core genome SNP alignment obtained by mapping sequence reads to the *P. aeruginosa* PAO1 reference genome (accession AE004091.2) (17). Gephi (45) was used to further interrogate interactions and visualise the effect size (observed/expected occurrences) of significant interactions (defined as Bonferroni-corrected binomial exact test *p*<0.01). In addition to effect size, significant interactions were also triaged by the frequency (n= 400 for DS only interactions and n= 300 for AGE interactions) of the AGE and their given D-value (D-value >0), a measure of phylogenetic independence that quantifies the phylogenetic signal of a binary trait. A lower D-value indicates a strong phylogenetic conservation (traits following Brownian motion evolution), while a higher D-value suggests more random distribution independent of phylogeny (46). To verify these findings with a more explicitly phylogenetically controlled method, Goldfinder was also used to identify interacting DSs and AGEs and detect co-occurrence and avoidance patterns between these elements (47).

**Figure 1.**
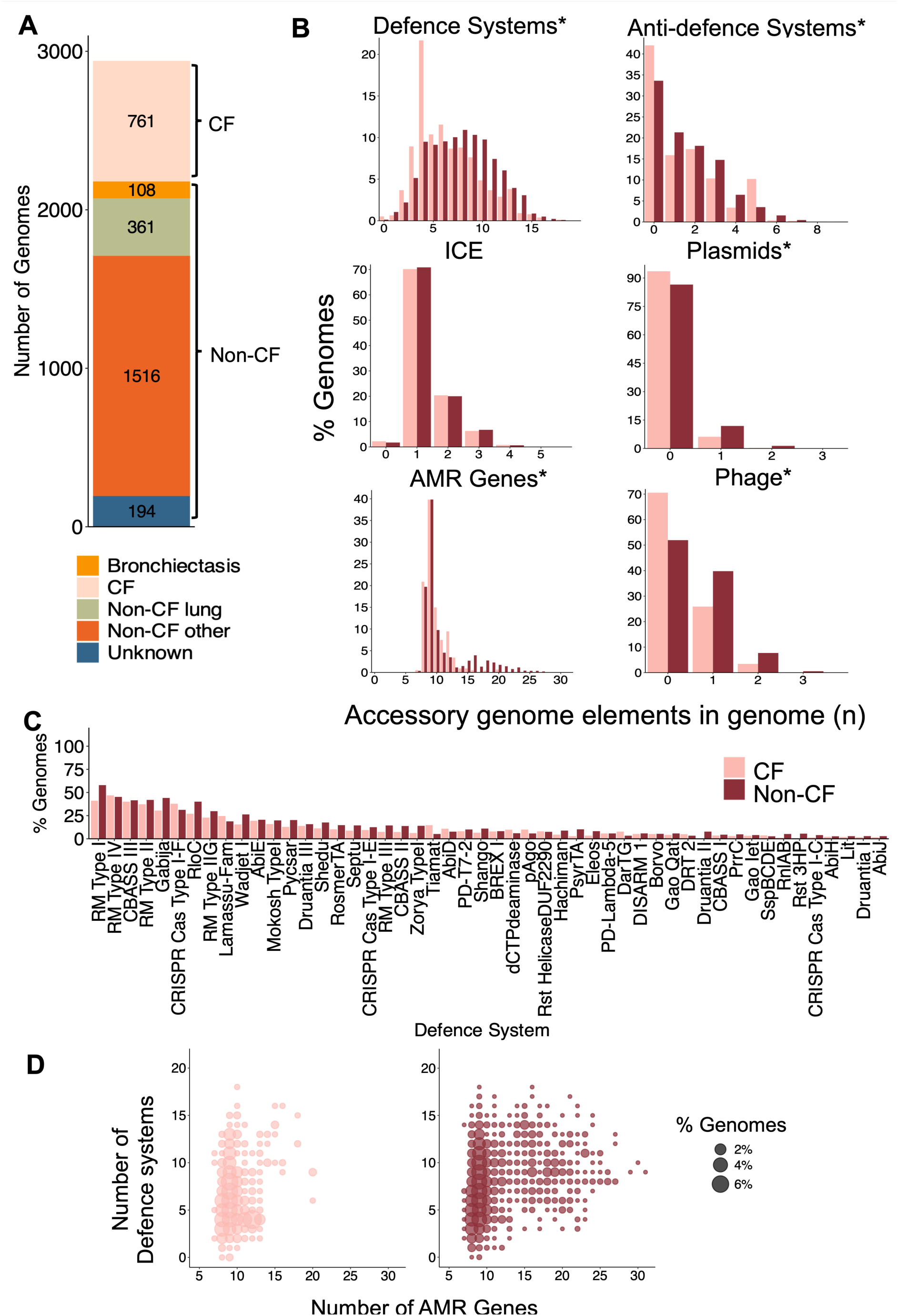
The distribution of accessory genome elements in *P. aeruginosa*. **(A)** Isolation niches for genomes in dataset (*n =* 2940. **(B)** The number of accessory genome elements (DSs, anti-DSs, ICEs, plasmids, ARGs and phages) per genome is shown as a percentage of genomes containing that number of element, stratified by CF/non-CF niche. Significant differences between the percentage of isolates containing any number of accessory genome elements from CF and non-CF niche’s is denoted with a * ((Wilcoxon rank sum *p*<0.05). **(C)** The percentage of genomes containing the 50 most prevalent DSs by niche (see Supplementary Table 1 for full dataset). ICE = Integrated Conjugative Element ARGs = Antimicrobial Resistance Genes CF = cystic fibrosis DS = Defence System. **(D)** Bubble plots showing the relationship between the number of AMR genes and defence systems per genome, where each point represents each genomic profile and bubble size reflects percentage of genomes with that profile, stratified by CF and non-CF niches.

### Statistical analyses

A Wilcoxon rank sum test was used to compare the frequency of AGEs in genomes between niches. The association of DSs with CF and non-CF genomes was tested using Fisher’s exact test with Bonferroni-adjusted p-values. T-tests were used to evaluate the differences in number of DSs, anti-defence systems and other AGEs per genome length (kb) between genomes isolates between niches, and to assess differences in DS frequency between isolates containing CRISPR-Cas Type I-F compared to those without. Effect sizes for t-tests were calculated using Cohen’s d, and for Wilcoxon rank sum tests using rank-biserial correlation. To assess the relationship between DS abundance and ARG content, we performed a simple linear regression analysis separately for both CF and non-CF isolates. To compare the length of contiguous sequences (kb) containing Pycsar and Shedu DSs (to those containing any other DS), we performed a Kruksall-Wallis rank sum test followed by a Dunn’s post hoc test for pairwise comparisons. All analyses were performed using R v 4.4.0.

To assess the relative importance of interactions among different AGE features, we calculated the proportion of significant interactions as a fraction of total theoretically possible pairs (among detected elements in the AGE types). All significantly interacting pairs were calculated based on the total number of each genomic element found (i.e. ARG = 441, anti-defence system = 30, DS = 130, ICE = 19, phage = 29, plasmid = 13 see Supplementary Table 1). To determine the total number of theoretically possible pairs between different AGE databases (i.e. types of accessory genome feature e.g. ARGs, DSs), we multiplied the size of the respective database (i.e. N_AMR_ x N_DS_, where *N* is the number of elements detected in the dataset). For pairs within the same category (e.g., AMR-AMR, we calculated the possible pairs as N_AMR_ x (N_AMR_-1)/2 to adjust for duplicates. The proportion of the observed number of significant interactions for each pair type was then divided by the total theoretically possible pairs for that category to determine the proportion of (possible) pairs that were significantly interacting.

## Results

### The accessory genome features of *P. aeruginosa* differ between niches

To explore the variation in the DSs of *P. aeruginosa* we analysed the DS repertoire of 2,940 *P. aeruginosa* genomes which were isolated from several sources (Figure 1A) and included globally distributed human, animal and environmental isolates. We grouped isolates into two categories: CF and non-CF and compared their frequency of AGE carriage (i.e. presence and absence). This division reflects two distinct ecological niches of *P. aeruginosa*, where CF isolates typically originate from chronically infected airways, with altered mucus composition, persistent infection, reduced microbial competition, repeated, but likely purifying antibiotic exposure, and strong host-derived selective pressures. In contrast, non-CF isolates are in this case associated with acute/transient infection, environmental reservoirs and transmission chains where exposure to more diverse competitors and MGEs is greater (48, 49).

This categorisation was further motivated by findings from the original study (Weimann, et al (2024)) (17) which reported a significant depletion of genes involved in bacterial defence in epidemic clones of *P. aeruginosa.* Therefore, we sought to examine whether similar patterns were evident in the distribution across a representative dataset of CF and non-CF isolates. Overall, *P. aeruginosa* contained an average of 7.5 DSs, with 99.8 % (n=2934/2940) of isolates containing at least 1 DS, consistent with previous studies (6, 18) (Figure 1B, 1C, Supplementary Table 2). Numerous DSs (n=130) were found, with RM Type I being the frequently found system, present in 53.6 % (n=1577/2940) of genomes. This is consistent with published observations stating that RM systems are present in >74% of prokaryotic genomes, with an average of ≥2 RM systems per genome (4). This also revealed that *P. aeruginosa* were rich in AGEs across both niches (i.e. DS (CF: 99.5% (n=757/761), non-CF: 99.9% (n=2176/2179)), anti-defence system (CF: 58.0% (n=441/761), non-CF: 66.4% (n=1446/2179)), plasmid (CF: 6.4% (n=49/761), non-CF: 13.4% (n=293/2179)), ARG (CF: 100% (n=761/761), non-CF: 100% (n=2179/2179)), ICE (CF: 97.8% (n=744/761), non-CF: 98.3% (n=2141/2179)) and phage (CF: 29.4% (n=224/761), non-CF: 48.0% (n=1047/2179)) (Figure 1A, 1B, 1C). We found that CF genomes (relative to non-CF genomes) were depleted for DSs, anti-defence systems, ARGs, plasmids, and phages (although these were weak effect sizes, Table 2). However, we also found that CF genomes were significantly smaller than non-CF genomes (Table 2) congruent with previous studies showing genomic streamlining in bacteria adapting to chronically infected lungs (compared with more acute infections) (17), and broader trends in host-adapting pathogens (50–55). However, our CF-associated AGE depletions remained statistically significant when frequencies were adjusted for genome size (t-test, *p*<0.05, Supplementary Figure 1).

**Table 2.** Summary of differences between genomes from CF and non-CF patients.

| Feature | Niche | Mean | Range (Min-Max) | IQR (Q1-Q3) | $p$-value | Effect size |
| --- | --- | --- | --- | --- | --- | --- |
| Genome Size (Mb) | CF | 6.5 | 5.9-7.5 | 6.3-6.7 | $9.18 \times 10^{55}$ | -0.64 |
|  | non-CF | 6.7 | 5.8-7.7 | 6.5-6.9 |  |  |
| DSs | CF | 6.5 | 0-18 | 4-9 | $2.14 \times 10^{25}$ | -0.25 |
|  | non-CF | 7.9 | 0-18 | 5-10 |  |  |
| Anti-defence Systems | CF | 1.5 | 0-7 | 0-2 | 0.017 | -0.05 |
|  | non-CF | 1.6 | 0-9 | 0-3 |  |  |
| ARGs | CF | 9.7 | 7-20 | 9-10 | $5.12 \times 10^4$ | -0.08 |
|  | non-CF | 11 | 7-31 | 9-12 |  |  |
| Plasmids | CF | 0.07 | 0-2 | 0-0 | $1.65 \times 10^7$ | -0.07 |
|  | non-CF | 0.2 | 0-3 | 0-0 |  |  |
| ICEs | CF | 1.3 | 0-5 | 1-2 | 0.92 | -0.001 |
|  | non-CF | 1.3 | 0-4 | 1-2 |  |  |
| Phages | CF | 0.3 | 0-3 | 0-1 | $1.16 \times 10^{19}$ | -0.19 |
|  | non-CF | 0.6 | 0-3 | 0-1 |  |  |

† Statistical tests performed: *t*-test for differences in genome size (Cohen's *d* effect size reported), *Wilcoxon rank-sum test* for all other features (rank-biserial correlation effect size reported).

To explore whether any individual DSs were enriched in either niche, we compared the proportion of genomes containing each DS per niche, revealing that 17% (n=22/130) of DSs were significantly associated with niche; 82% (n=18/22) with the non-CF niche and the remainder with the CF niche (Bonferroni-corrected Fisher’s Exact Test, *p*< 0.05). The DSs associated most significantly with non-CF and CF niches were Druantia II and Tiamat respectively (Figure 1C). RM Type IV was found in the most genomes (n=357/761) within the CF niche, while RM Type I was the most common (n=1264/2179) in non-CF isolates.

And finally, to determine whether the ecological differences between CF and non-CF environments influence accessory genome composition, we compared the number of DS and ARGs across isolates from each niche. Consistent with the global comparison results (Table 2), there were a higher number of ARGs in non-CF isolates, though the distributions highlight a long tail of genomes with a high number of ARGs (Figure 1D). This is consistent with isolates in the non-CF environment retaining a higher number of AMR genes for survival in the harsher, more diverse environment and suggests that DS and ARG repertoires are shaped, in part, by niche-specific selection pressures. Despite this influence however, the overall effect sizes of the CF/non-CF niche was weak (Table 2), and as other studies have suggested that the distribution of AGEs is shaped by the ecological (i.e. cooperative vs antagonistic) relationships among DSs (and other AGE classes, Table 1) themselves. We then progressed to conduct a global analysis of the data to explore the potential interactions among the diverse AGEs in *P. aeruginosa* with a view to identifying putative associations and disassociations of DSs (and other AGEs) that might represent putative mechanistic relationships (while controlling for phylogenetic relationships).

### Determining defence system interactions

To identify putative synergistic or antagonistic DS interactions across *P. aeruginosa*, we explored which DSs were more frequently associated (putative synergism) or dissociated (putative antagonism) with each other than expected by chance and independent of evolutionary relationships. For this we first, employed CoinFinder (43), which identifies associations and dissociations between traits (in our case, accessory genome categories of elements) and also estimates the phylogenetic independence (as measured by D-value) of traits.

Association analysis with Coinfinder revealed 182 pairwise interactions among DSs (27.9%, n=182 of 653 possible pairs) that were significantly associating or dissociating with one another (Bonferroni-corrected binomial exact test *p*<0.01, Figure 2, Supplementary Table 3 & 4). Association interactions made up 85.2% (n=155/182) of the total significant DS pairs and the remaining interactions were dissociations. Overall, the dominance of associations over dissociations supports current hypotheses that DS complement each other (14, 18, 56, 57). Each interacting DS (n=36) had a variable number of associations and dissociations, with an average of 10 interactions (range 2-19), and RM Type I having the most interactions (n=19). The high frequency of interactions between RM Type I and other systems may reflect that RM shapes the DS complement of genomes. Other DSs also exhibited high numbers of associations, such as RM Type IV, Pycsar and AbiE, which all associated with 17 other DSs (Figure 1C, 2 & Supplementary Table 3 & 4).

**Figure 2.**
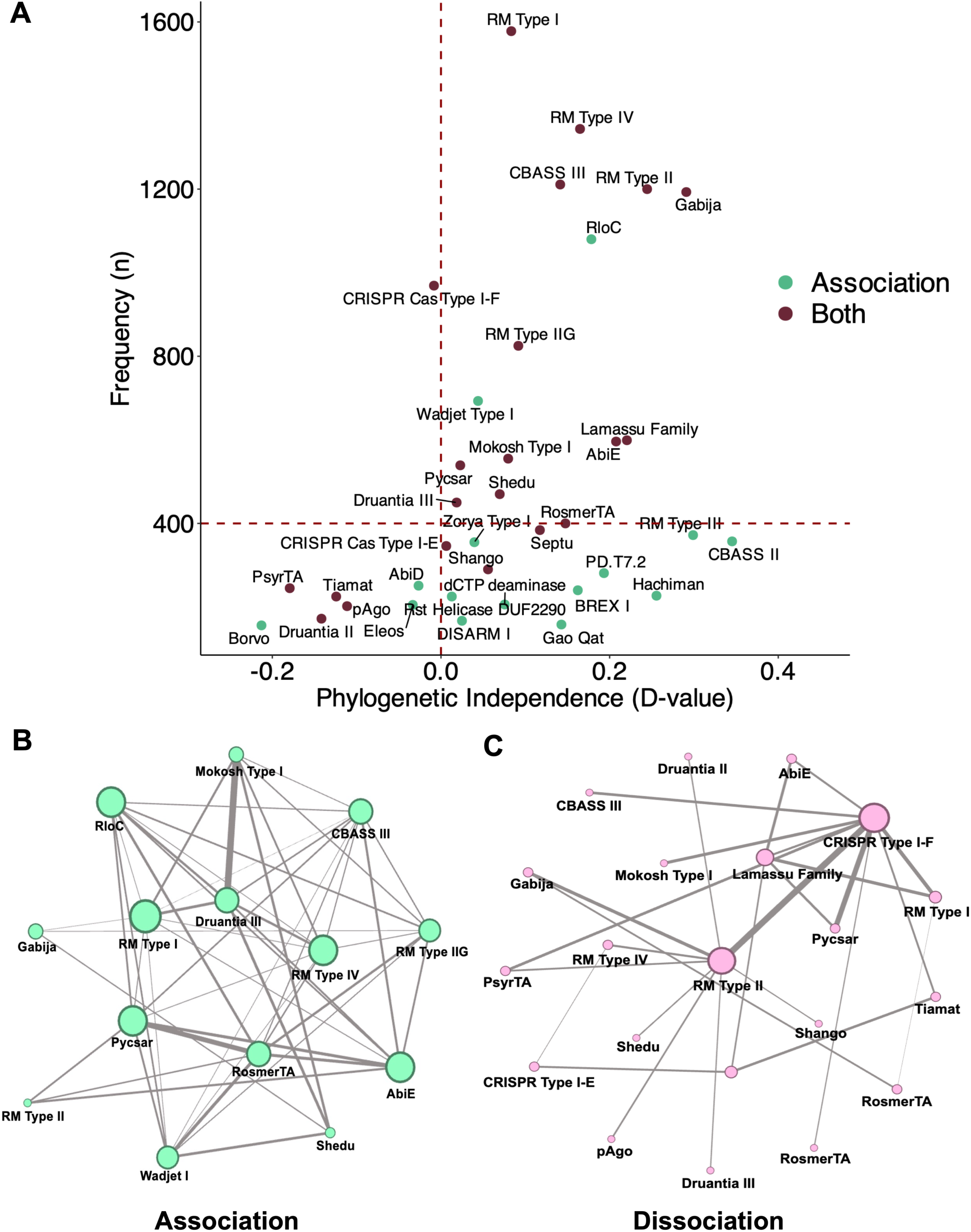
The distributions, associations and dissociations of defence systems in the global *P. aeruginosa* population. **(A)** The frequency and phylogenetic independence of DSs with significant interactions (i.e. associations or dissociations). Green dots highlight DSs that were found to have significant associations while purple dots indicate DSs that had both significant associations and dissociations (no systems were found to only have significant dissociations, full data in Supplementary Table 3 & 4). Red dashed lines show triage cut-off values of 0 and 400 for Phylogenetic independence and Frequency respectively. **(B)** Interaction network of associations among DSs. Each node depicts a DS scaled in size by the number of significant associations it had. Edges represent individual associations between DSs with the thickness corresponding to the effect size (i.e. strength of association). **(C)** Interaction network of DS dissociations. Each node similarly depicts a DS scaled in size by the number of significant dissociations it had. Edges represent individual dissociations between DSs with thickness corresponding to effect size.

To focus our attention on those associations that may have significant roles in shaping *P. aeruginosa* populations, we triaged DSs to focus on those that were comparatively mobile and more common. As a proxy for mobility, we used the phylogenetic independence (D-value) of each DS. As a lower D-value indicates a strong phylogenetic conservation we considered DSs with a D-value of over 0 (Figure 2A) in our downstream analysis. However, we note that the overall D-values across traits were relatively low (ranging from 0.02-0.4), suggesting that while these DSs were not fully phylogenetically conserved, some dependence on phylogeny likely remains. This limitation is considered in our interpretation of the results. Additionally, we filtered the interactions to only include DSs that were present in over 400 (13.6%, n= 400/2940 of the total dataset, full data in Supplementary Tables 3 & 4) genomes (Figure 2A). Of the 15 DSs that passed these thresholds, 13 were found to both associate and dissociate with other systems, with the remaining 2 having only associations (Figure 2A). Among these 15 DSs, RM Type II, Gabija Lamassu and AbiE were the most frequent and exhibited the more phylogenetic independence (higher D-value within our range) (Figure 2A).

Having triaged on frequency and phylogenetic independence, we then sought to interrogate DS interactions by visualising the effect size (inferred from the ratio of observed vs expected occurrences, methods) of each association alongside the number of interactions of each DS (Figure 2B & C, Supplementary Table 3 & 4). This revealed that among the 54 (n= 54/155 total associations) interactions between the 15 triaged DSs, Druantia III and Mokosh Type I association exhibited the strongest interaction (Figure 2B). This indicates a possible mechanistic synergism and is consistent with experimental evidence suggesting Druantia III and Mokosh Type I both protect against similar phages such as T4, through nuclease-based mechanisms (7, 58, 59).

We then examined DS pairs that were found to dissociate from each other. As these only made up only 14.8% (n=27/182) of the total significant interactions (Figure 2C & Supplementary Table 4) we applied no triage thresholds (though notably those with a D-value of <1 should be viewed with caution owing to the potential phylogenetic association of individual DSs, see Figures 1A & C). Here we found that CRISPR-Cas Type I-F had the most dissociations with other DSs, such as other RMs Type I & II and Pycsar (Figure 2C & Supplementary Figure 2). Isolates containing CRISPR-Cas Type I-F (n=969/2940) contained fewer DSs, compared to isolates not containing CRISPR-Cas Type I-F (t-test, *p*<0.05) (Supplementary Figure 2). These strong indicators for dissociation of CRISPR-Cas Type I-F are consistent with it being commonly found in *P. aeruginosa* and blocking HGT of foreign DNA and even other DS (5, 60, 61). To better control for phylogenetic dependence, we also ran the program Goldfinder to detect interactions between DSs, complementing the Coinfinder analysis. This additional approach was taken because the D-values for each significantly interreacting DS were relatively low, indicating that many traits showed some phylogenetic association. Encouragingly, Goldfinder identified 9 associations among 11 DSs (with D-values of between 0.02 and 0.4, Supplementary Table 3 & 6), most of which (66.7%, n=6/9 pairs) were also detected by Coinfinder and were within our filtering thresholds.

As recent studies have shown that DSs are frequently encoded within horizontally transferrable defence islands or genomic hotspots (3, 9–11, 58) we sought to explore whether our DS associations resulted from co-localisation. Therefore, we assessed whether the significant associating and dissociating DS pairs (n= 77, Figure 2B & C) were present on the same contiguous sequence (contig) in our draft genome assemblies (as an imperfect but practicable proxy of co-localisation, Figure 3). This revealed that of all DS containing contigs, only 29% (n=7086/24211) carried two or more DSs. Within the 7086 contigs containing ≥2 DS, associating DS pairs were co-located in only 22% of cases i.e. were on the same contig 4,367 times out of a total of 19,472 pairs occurring across genomes (Figure 3). Notably, the six intersecting associations detected by Goldfinder had some of the highest (>0.5 proportion) rates of co-localisation (Figure 3, Supplementary Table 6). As a counterfactual for understanding the rates of co-localisation among associating DS pairs, we examined co-localisation rates among dissociating DS pairs (Figure 3). As anticipated, the rates of co-localisation were far lower i.e. 0.5%, n=175/37,098 pairs across all genomes, though this varied among interacting pairs (Figure 3). We found the rate of co-localisation for 22 (of 54) associating pairs was lower than 0.034% (with 0.034% being the highest co-localisation proportion for a dissociating pairs) (Figure 3). We observed that some associating DSs were frequently below this 0.034% threshold (e.g. Pycsar) potentially indicating genuine ecological interactions not from co-localisation, and we further verified that this was not attributable to an overall shorter length of contigs containing those DSs (Figure 3 & Supplementary Figure 3). Overall, our results indicate that many associations detected here are unlikely to have entirely resulted from co-localisation.

**Figure 3.**
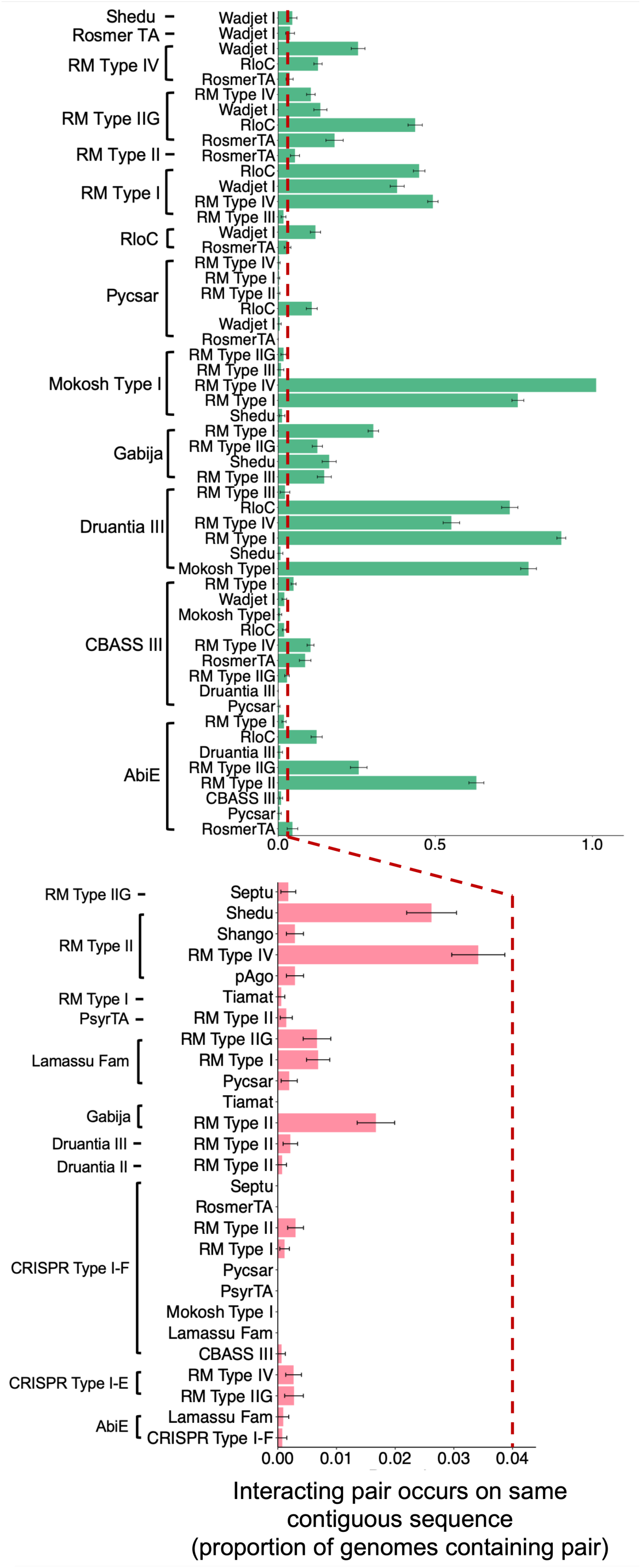
The proportion of pairs of DSs showing interactions of association (upper, green) and dissociation (lower, pink) that occur on the same contiguous sequences in draft assemblies. Axes labels are ordered alphabetically. Error bars represent standard error. The red dashed line intersects the x-axis at 0.04 on both plots to highlight the difference in scale and show that some associating pairs were less frequently co-located than dissociating pairs.

Hence, our analyses of DS interactions in *P. aeruginosa* revealed a complex network of both association and dissociation interactions. Most significant interactions were associations, with DSs such as RM Type II, Gabija, Lamassu and AbiE frequently associating with other systems. Strong interactions between DSs may reflect complementary functions. Conversely, frequent dissociations with CRISPR-Cas Type I-F indicates its role in restricting HGT and conferring a selective disadvantage to co-occurring DSs. Although co-localisation of DSs on contigs was generally low, some associations such as Mokosh Type I and RM Type IV were consistently (n=415/415 genomes containing the pair) co-located. While such co-localisation can suggest potential functional synergy (57, 62), in this case, the less-phylogenetically dependent co-occurrence patterns could also indicate that co-inheritance could be driven by genetic linkage rather than necessarily reflecting synergistic interactions between the systems (Figure 2A). Environmental context may also contribute to these patterns, as DS content can be shaped by local ecological conditions or exposure to specific phage communities. in addition to interactions with plausible functional explanations, our analysis also identified numerous statistically significant DS interactions for which no biological link has yet been described (e.g. CBASS II-Wadjet I & Gabija-Tiamat). These novel patterns may represent unexplored functional compatibility or incompatibility between DSs and therefore warrant further experimental investigation. Overall, our findings highlight the dynamic interplay between DSs, influenced by both genomic context and environment, that may shape the adaptive potential of *P. aeruginosa* populations.

### Interactions of DS with other accessory genome elements

To further explore the complexity of integral links among DSs and other AGEs, including the recently described anti-defence systems, we used similar methods to characterise the interactions among DSs and all other AGEs that maybe influencing accessory genome composition. Coinfinder analysis revealed a total of 591 significant interactions between specific AGEs distributed across accessory genome feature categories (Figure 4 & Supplementary Table 3 & 5). As with DS comparisons, we triaged the interactions by frequency (using those present in 7%, n=300/4288) of each element across the dataset and phylogenetic independence (D-value > 0) (Figure 4A). Analysis of the 36 (of 68) AGEs meeting these triage criteria revealed a network with 201 associations of various effect sizes and differences in the number of associations (ranging between 2 and 31 per element) (Figure 4A & Supplementary Table 3 & 5). Of the 201 pairs passing the threshold, anti-CRISPR AcrIF3 and anti-CRISPR AcrIF5 show the strongest association (Figure 4A & Supplementary Table 5). Other comparatively strong associations included those between ARGs *qacE*Δ and *sul1*, and DS AbiE and ICE bhp-sal (Figure 4A & Supplementary Table 5).

**Figure 4.**
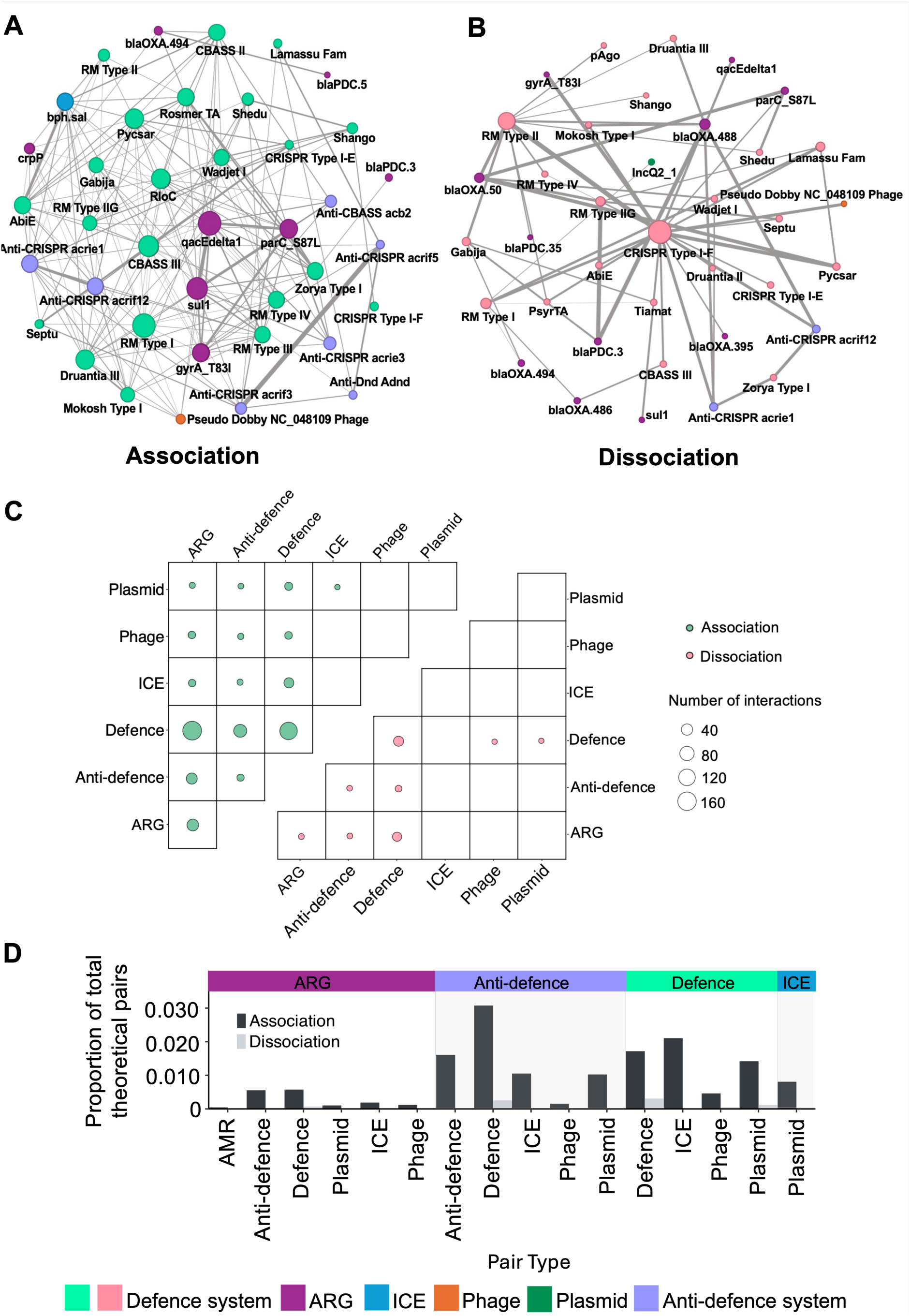
Interactions among accessory genome elements in *P. aeruginosa*. **(A)** Significant associations network among individual accessory genome elements. Each node depicts a genomic element coloured by type according to the inlaid key. Nodes are scaled by the number of significant associations the element has. Edges represent significant associations between elements with thickness scaled by effect size. Elements have been triaged using a Phylogenetic independence value of ≥0 and being found in >300 genomes. **(B)** Significant dissociations network among individual accessory genome elements. Each node depicts a genomic element coloured by type according to the inlaid key. Nodes are scaled by the number of significant dissociations the element has. Edges represent significant dissociations between elements with thickness scaled by effect size. **(C)** Bubble plots representing significant associations (upper) or dissociations (lower) interactions between pairs of accessory genome feature categories (axes). Bubbles are scaled by the number of significant interactions between those categories. **(D)** The relative importance of AGE interactions is shown as bar plots showing the proportion of feature category pair interactions that were found to be significant associations (dark grey) or dissociations (light grey) with the denominator of the total number of theoretical pairs (to account for database size). Feature category pairs are indicated by groups above the bar plots (coloured according to the inlaid key) and x-axis labels.

Regarding the dissociation interactions among these individual AGEs, CRISPR-Cas Type I-F was again found to have the most negative interactions (29.8%, n=17/57 total dissociations with no triage criteria applied). The significant dissociation between CRISPR-Cas Type I-F and RM Type II was also exhibited in this analysis, further supporting the observation that they may not act synergistically in *P. aeruginosa* (Figure 4B & Supplementary Table 5). The opposite is seen for the Gram-positive bacteria *Listeria*, *Streptococcus* and *Staphylococcus*, where CRISPR-Cas Type IV and VI systems have been observed to provide synergistic protection against phage infection with RM systems suggesting synergy maybe system type specific and species specific (63–65). We also observed that the ARG blaPDC3 had two strong dissociations with both RM Type IIG and blaOXA-488 (Figure 4B & Supplementary Table 5). Notably, we also observed distinct patterns of interaction between DSs and their cognate inhibitors, where anti-CRISPR systems associate with CRISPR-Cas Type I-E but dissociate from CRISPR-Cas Type I-F and anti-CBASS systems associate with CBASS systems.

Although we again applied a D-value threshold of >0 to reduce phylogenetic dependence, the overall D-vales were relatively low (highest = 0.5), indicating that some phylogenetic structure may remain in these data. To further evaluate these associations, we re-analysed the dataset using Goldfinder, which more explicitly controls for phylogeny. Goldfinder identified 14 of the 36 filtered AGEs as positively interacting with each other and detected 19 significant associations among them (with D-values of between 0.02 and 0.4, Supplementary Table 3 & 7). Of these pairs, 11 were also identified by Coinfinder, indicating some concordance between the two methods. No dissociations were found by Goldfinder, and notably, in line with our AGE interaction networks (i.e. Figures 4A & B) being dominated by DSs and anti-defence systems, GoldFinder only found interactions among DSs and anti-defences. The recovery of mainly DS and anti-defence in our interactions supports this strategy as having detected putatively sub-genomic ecological interactions among AGEs and prompted us to further explore the broader trends among AGE groups in terms of quantitating interactions among AGEs.

To characterise the broader dynamics and relative importance of interactions among AGEs, we collapsed our AGEs into feature classes (i.e. ARGs, DS, phages, etc) according to the respective feature databases and correlated this to the detected association and dissociation interactions (with associations again predominating [90.4% n=534/591 interactions]). This revealed the highest numbers of associations between ARGs and DSs (31.1%, n=166/534 interacting pairs) followed by DSs and DSs (27% n=144/534) and anti-defence systems and DSs (n=11% 60/534, Figure 4C). Conversely, ARGs and DSs (35.1% n= 20/57) along with DSs and DSs (45.6% n=26/57) had the most dissociations (Figure 4C). However, these results do not account for discovery bias that may have arisen from differences in detection of AGEs attributable to the respective size of the databases. Hence, to use our interaction findings to bring meaningful insights on the relative importance of AGE interactions, we calculated how many interactions arose between AGE classes as a proportion of the total number of theoretical pairs that could occur between each pair of AGE classes (e.g. for ARG to DS interactions, there were a total of 166 significant associations among a theoretical total of 28,665 possible interactions, enumerated from the detection of 441 unique ARGs and 130 unique DS, giving a proportion of 0.006 significant pairs (Methods, Figure 4D). Overall, this revealed that DS association interactions with themselves and with anti-defence systems and ICEs were comparatively overrepresented (Figure 4D), and that self-interactions between anti-defence systems were also high. Dissociation interactions, although lower in abundance, were more frequently detected between DS-DS, anti-defence-DS, and plasmid-DS pairs (Figure 4D). Hence, our results indicate that interactions between DSs and anti-defence systems are common within the accessory genome and may influence bacterial genome composition. While such interactions are consistent with an ongoing interaction among DSs and anti-defence systems, they do not necessarily imply direct functional synergy, and these patterns will be further influenced by other ecological factors. Instead, they highlight the complex interplay of factors shaping genetic diversity, AMR and bacterial evolution (3, 66, 67).

## Discussion

In this study we investigated AGEs across ecological niches of global *P. aeruginosa* population to find biologically meaningful relationships among the densely packed accessory genome.

The comprehensive analysis of 2,940 *P. aeruginosa* genomes revealed 82% of detected DSs were significantly associated with bacterial niche, highlighting the role of environmental pressures in shaping genome defence strategies. Notably, the CF niche isolates were enriched for DarTG and CRISPR-Cas Type I-F, while non-CF niche isolates exhibited a higher prevalence of Druantia II and RM-Type I systems. These findings suggest that CF-adapted *P. aeruginosa* may rely on distinct defence mechanisms, potentially reflecting selection for systems that counteract host-adapted phages or environmental stress (respectively). Different bacterial DSs may be favoured under distinct ecological conditions, yet identifying the environmental drivers of their distribution in complex environments remains challenging. Although, there are studies that have previously linked some ecological factors such as force of infection to the evolution of immune strategies, where bacteria in lower infection risk environments can favour CRISPR immunity compared to surface modification immunity (68, 69). This is supported by increased abundance of CRISPR systems in thermophiles that reside in low infection risk environments (70, 71), which may partly explain the enrichment of CRISPR-Cas Type I-F in *P. aeruginosa* isolates from the CF niche in this study. Furthermore, pan-genome differences between different *P. aeruginosa* phylogroups suggest that variation in DSs could influence homologous recombination dynamics. However, these associations may also reflect ecological or phylogenetic factors rather than direct functional effects. While more diverse or abundant systems might limit the uptake of exogenous DNA, this remains a hypothesis requiring further validation (72). Additionally, CF isolates exhibited significant genome reduction and contained fewer DSs, anti - defence systems, plasmids, ARGs and phages per kilobase of genome length, supporting the hypothesis that long-term adaptation to the host-lung environment drives genomic streamlining, through loss of genes linked to motility, virulence and environmental adaptability due to focused selection pressures in chronic infection. However, as these were overall weak effects, we sought to determine whether ecological relationships among the sub-compartments of the vast accessory genome could be driving genome composition.

We found a high density of DSs in *P. aeruginosa* genomes, consistent with previous studies, and extended this work by illuminating the many significant interactions among DSs. Associating interactions were far more common than dissociating ones, supporting the hypotheses that DSs function in a complementary manner, working alongside each other to provide a multi-layered approach against invading AGEs, providing protection if other defence mechanisms fail (6, 14, 18, 56–58, 73, 74). Distinctly however, we reveal that associations are unlikely due to co-localisations in defence islands.

Conversely, CRISPR-Cas Type I-F had the highest number of dissociations, particularly with RM systems and Pycsar. Given its well documented role in blocking HGT, including the acquisition of other DSs (60, 61, 75), this may explain why genomes containing CRISPR-Cas Type I-F harboured significantly fewer DSs than those without it (Supplementary Figure 2). Together, these findings highlight the complex interactions shaping DS organisation and suggests that while many DSs complement one another, CRISPR-Cas Type I-F may act as a gatekeeper, limiting genomic plasticity in *P. aeruginosa* living in potentially nutrient deficient, highly competitive or stressful environments. Hence, the interactions we show among defence systems here represent plausible meaningful mechanistic or ecological relationships beyond simple co-inheritance on defence islands. While these patterns are consistent with potential interactions, they do not necessarily impact direct functional synergy and could also arise from shared environmental or phylogenetic factors (7, 58, 76).

Overall, our study of the accessory genome interactions in *P. aeruginosa* revealed a complex network of interactions among AGE classes that will be partly shaped by ecological interactions. However, we also found relationships among AGEs that were consistent with ecological interactions among AGEs (e.g. direct sub-genomic conflicts) shaping the distribution of AGEs. For example, the most frequent interactions were between ARGs and DSs, consistent with anti-plasmid activity, followed by DS-DS interactions and DS-anti-defence system interactions, consistent with putative mechanistic interactions among these AGEs and suggesting that DSs play a central role in structuring the *P. aeruginosa* accessory genome. Many defence genes are found in the accessory genome and are a key contributor to intraspecific variation of bacteria (77, 78). However, their high turnover rate, co-acquisition with AGEs and persistence in the accessory genome all affect the amount of protection that they offer against HGT (69, 79, 80), as will shifting environments, as highlighted above.

At a finer scale, our data revealed strong associations between specific AGEs, such as anti-CRISPR AcrIF3 and AcrIF5, as well as the ARG pair *qacE*Δ and *sul1* and the DS AbiE with ICE bhp-sal. These findings reinforce the notion that AGEs interact to promote dissemination and survival in the environment, where AcrIF5 works closely with AcrIF3 to block Cas 2/3 recruitment to ensure that CRISPR-Cas Type I-F cannot degrade foreign DNA (73). The spread of ARGs is facilitated somewhat by DSs, where systems such as Abi, Gabija, CBASS and RM have been found to be more frequently located within or next to AGEs like ICEs, to affect AGE propagation, stability and HGT to confer resistance in bacteria (3, 58, 79, 81, 82). Our study reveals key associations and dissociations that appear largely independent of phylogeny (though this should be explicitly considered alongside the results here e.g. D-values, the respective findings between Coinfinder and Goldfinder). These patterns may be further shaped by multiple factors including ecological context, and differential selective pressures.

Which brings us to the limitations of this study. Firstly, although our dataset includes isolates from diverse ecological niches, many isolates originate from studies on AMR, which may bias genome composition. Our findings are also reliant on the content and curation of various AGE databases, so may have failed to detect significant interactions with yet undescribed AGEs. Furthermore, our association interactions appeared to be robust against incidental co-localisation but ascertaining the full impact of co-localisation would be better evaluated with complete genomes assembled from long-read sequencing data (e.g. as in Wu et al., 2024). Another limitation is that we relied solely on DefenseFinder (6) to identify DSs, which may not fully capture the diversity of all DSs. As a result, we may have missed relevant interactions due to incomplete or imperfect annotations. A further consideration is that despite trying to account for phylogenetic independence using the Coinfinder D-value output, the overall values were relatively low, indicating that some phylogenetic dependence likely remains. To address this, we re-analysed our data using Goldfinder (47), which accounts for shared ancestry by mapping trait distributions onto the phylogeny and identifying interactions that are independent of evolutionary relatedness. Despite identifying fewer total interactions, Goldfinder recovered many of the same associations between DSs and AGEs as Coinfinder, supporting the robustness of our main findings although these were often highly co-located, so each interacting pair should be interpreted alongside markers of co-localisation and phylogenetic independence. Our *in-silico* investigation into putative synergism or antagonism among AGEs in *P. aeruginosa* does not offer explanations of the mechanistic or ecological basis for those interactions. However, the approach is robust to many confounding factors (e.g. phylogenetic dependence and the impact of genome size and co-localisation), and many had a plausible mechanistic basis, so we put forward these interactions as strong candidates for future research on biologically meaningful interactions among the accessory genome milieu in *P. aeruginosa*.

The results of this work highlight the importance and complexity of accessory genome interactions in bacteria, particularly the expanding array of DSs and anti-defence systems in shaping accessory genome interactions. Multiple association and dissociation interactions between DSs, and other accessory genome features were found and suggest adaptation of *P. aeruginosa* is driven by the selective forces in their environment. Future research should aim to explore and validate these putative interactions both phenotypically and mechanistically, and to continue to explore other possible drivers of this apparent synergy and antagonism.

## Supplementary Material

**Supplementary Table 1**. *Pseudomonas aeruginosa* isolates, associated metadata and accessory genome feature content used in this study.

**Supplementary Table 2.** DefenseFinder and Anti-defenseFinder output from *P. aeruginosa* dataset.

**Supplementary Table 3.** Accessory genome elements found to significantly interact by Coinfinder, their associated D-value, frequencies and interactions.

**Supplementary Table 4**. List of DSs pairs found to associate and dissociate by Coinfinder.

**Supplementary Table 5**. List of accessory genome feature pairs found to associate and dissociate by Coinfinder.

**Supplementary Table 6.** List of DS pairs found to associate and dissociate by Goldfinder.

**Supplementary Table 7.** List of accessory genome feature pairs found to associate and dissociate by Goldfinder.

**Supplementary Figure 1.**
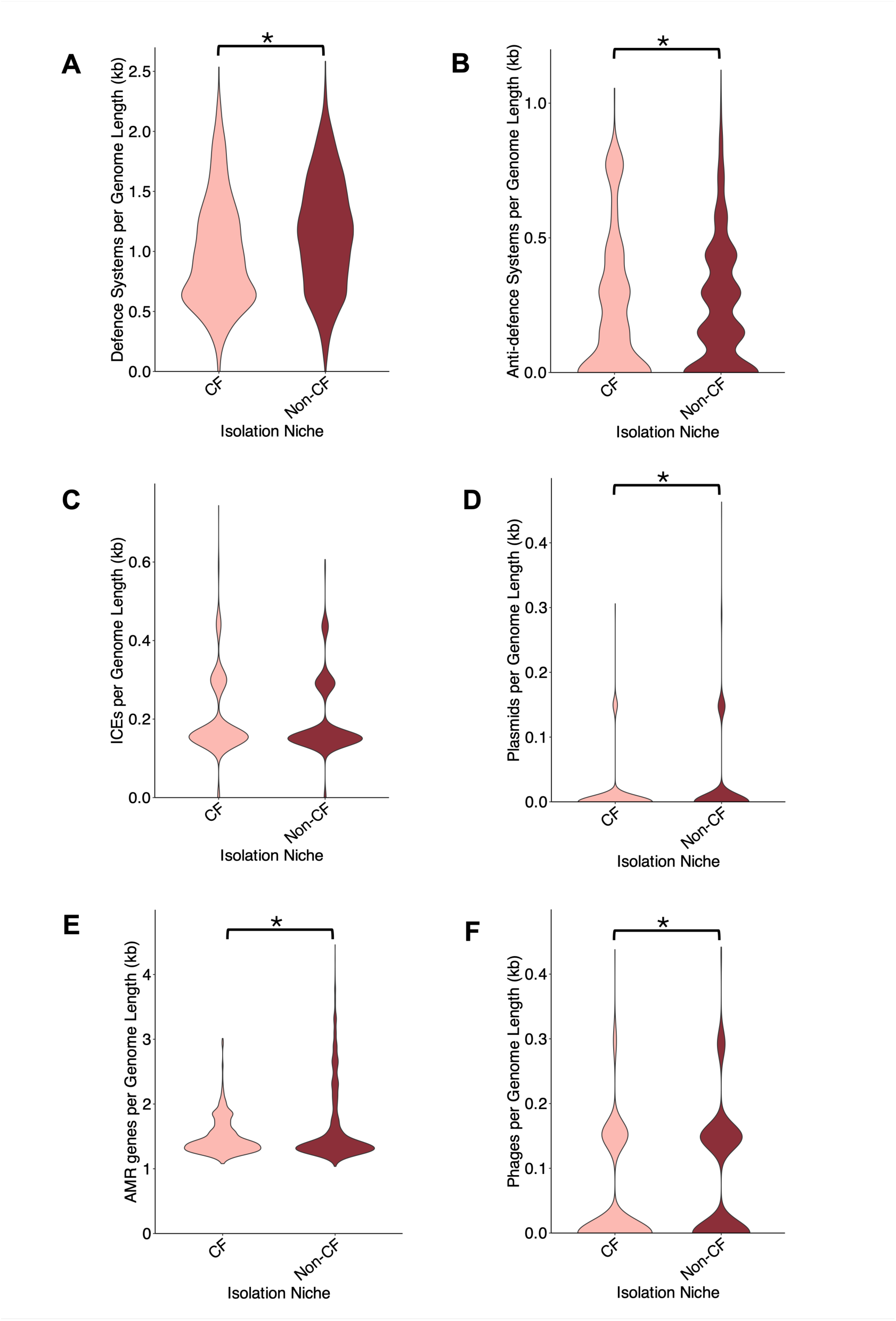
Number of defence systems **(A)**, anti-defence systems **(B)**, ICEs **(C)**, plasmids **(D)**, ARGs **(E)** and phage **(F)** present in *P. aeruginosa* per genome length (kb) of genomes isolated from CF patients and non-CF sources. Significant differences in number of accessory genome elements per genome length (kb) between genomes isolates from CF and non-CF niches is denoted with a * (*t*-test *p*<0.05). ICE = Integrated Conjugative Element ARGs = Antimicrobial Resistance Genes CF = cystic fibrosis DS = Defence System.

**Supplementary Figure 2.**
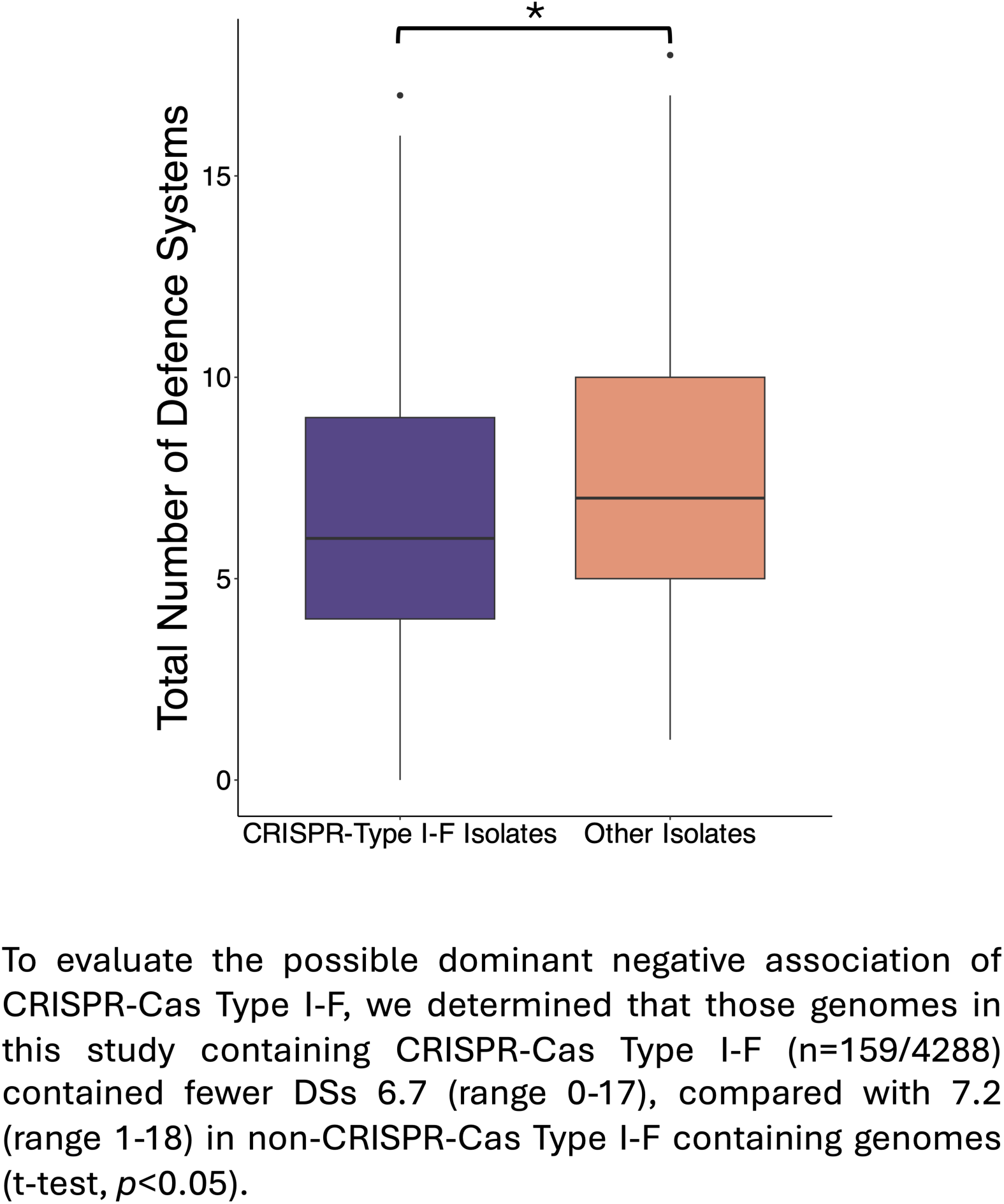
Comparison of the number of DSs found in *P. aeruginosa* isolates containing the DS CRISPR-Type I-F (n=1591) and isolates that did not (n=2683). The plot shows the median (line), interquartile ranges (box), and range (whiskers) of the number for each group. Outliers are shown as individual markers. The asterisk indicates a statistically significant difference *(p*<0.05) as measured by a *t*-test. Shedu and Pycsar, which were on average found to co-localise in lower proportions, compared to other non-Pycsar/Shedu containing pairs (average proportion of co-localisation in DS pairs containing, Pycsar = 0.01, Shedu = 0.06 and all other system pairs= 0.2. To confirm that these apparently low-rates of non-co-localisation of Pycsar and Shedu were not the result of disproportionately shorter contigs, we assessed the average contig length (kb) of sequences containing these DSs relative to other OS-containing contigs. This revealed that Shedu or Pycsar containing contigs (average of 393 kb and 349 kb respectively) were slightly longer than contigs containing other DSs (average of 275 kb) (Kruksall-Wallis rank sum test and Dunn’s post hoc test = *p<0.05)* suggesting that this is a genuine interaction between Shedu and Pycsar, and other systems and might represent either functional redundancy or mechanistic antagonism with other systems.

**Supplementary Figure 3.**
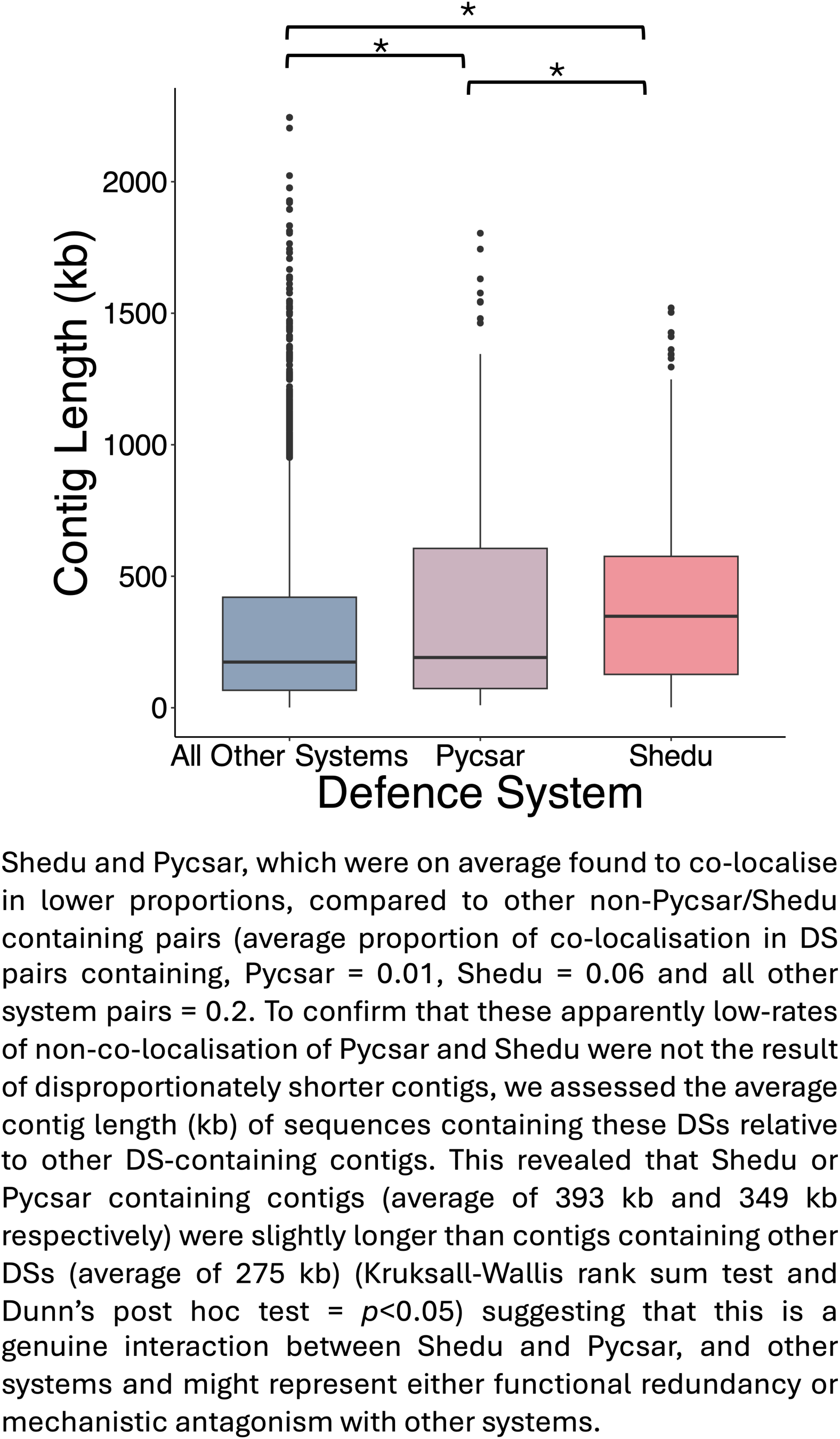
Comparison of length in kilobases of contiguous sequences containing either Pycsar (n=539), Shedu (n=586) or all other DS (n=23471). The plot shows the median (line), interquartile ranges (box), and range (whiskers) of the number for each group. Outliers are shown as individual markers. The asterisk indicates a statistically significant difference (*p*<0.05) as measured by a Kruksall-Wallis rank test followed by a Dunn’s post hoc test for pairwise comparisons.

## Author contributions

C.E.C and K.S.B contributed to the writing of the original draft. All authors (C.E.C, A.W, A.A, J.L.F, M.A.B, J.P, R.A.F, M.D.S, E.R.W, M.C and K.S.B) contributed to reviewing and editing of subsequent drafts. C.E.C was responsible for formal analysis, investigation, visualisation and methodology. A.W, J.P, R.A.F, and C.E.C. contributed to data curation. K.S.B, M.A.B, M.D.S, C.E.C, and E.R.W contributed to funding acquisition and conceptualisation. K.S.B and E.R.W were responsible for project administration. K.S.B was responsible for supervision.

## Acknowledgements

We thank all current and past members of the Multi-Defence Consortium for providing scientific support.

## Data availability

Short-read DNA was downloaded from the European Nucleotide Archive (ENA) by Weimann et al. (2024). ENA run accessions can be found in Supplementary Table 1.

